# Exploring Phylogenetic Classification and Further Applications of Codon Usage Frequencies

**DOI:** 10.1101/2022.07.20.500846

**Authors:** Logan Hallee

## Abstract

Codons are the repeats of three nucleic acids in genetic material read during translation. 64 total codons exist at different frequencies known to vary between organisms. Codon usage frequencies (CUFs) have recently been used for phylogenetic classification at different discrimination levels. However, the accuracy of phylogenetic classification and applications of this predictive power are not fully elucidated in the current literature. The goal of this project was two-fold: 1.) To increase the accuracy and depth of phylogenetic classification models using CUFs in literature. 2.) To explore the potential application of identifying open reading frames (ORFs) with CUFs. To increase the accuracy of classification models GridSearchCV, TensorFlow, and keras were used to design an improved Artificial Neural Network than the relevant example in the literature. Commonly used predictors were explored in an ensemble format that performs even better than the improved neural network. To explore a more discriminatory and phylogenetically deep classification K Means was used to look at possible clustering structures in the CUF data. To identify ORFs the codon frequencies of each potential ORF are compared to the frequencies of an organism of choice with a multinomial goodness-of-fit test. With correct optimization, these tests can reject possible ORFs with high confidence. In addition to these goals, the codons were ranked in terms of importance for classification with lasso regression and random forests feature ranking. This not only highlights exciting biology related to tRNA concentrations and the variance thereof, but is also helpful for optimizing the statistical tests for ORF identification.

## 1 Introduction

### 1.1 Project Description

The genetic code is what is known as redundant, composed of four molecules but read three at a time. This results in 4^3^ or 64 total three molecule combinations, known as codons. The frequencies of codon usage throughout an organism have emerged as a recent marker to classify DNA sequences phylogenetically [1]. However, this initial approach in current literature is insufficient in fully elucidating the possibilities of codon usage frequency (CUF) phylogenetic classification. Upon experimentation, the models presented here improve upon the classification by [1] with various kingdom structures. Even simple CUF classification models perform well at different levels of discrimination. For example, they nearly perfectly classify eukaryotes vs. prokaryotes and have high accuracy when classifying bacteria vs. viruses vs. animals vs. plants vs. archaea. So how good can the classification get? The first goal of this project is to improve the classification from [1] and explore how specific the classification can go. Perhaps, between eukaryotes vs. prokaryotes and species vs. species, there is a line of effective phylogenetic classification that CUFs can model. To explore this line K Means is used with a vast range of starting centroids to look at potential structures in the CUF data. If there is a high *k* structure in the data then it should be possible for the phylogenetic classification to be more discriminatory (biological class vs order for example).

However, these different phylogenetic categories can already be classified in many ways; why are CUFs worth exploring? One interesting potential application is in open reading frame (ORF) identification. Genome annotation is a notoriously tricky problem. Sequencing entire genomes is no longer a challenge with next-generation sequencing techniques that are gradually becoming inexpensive over time. Nevertheless, interpreting the billions of DNA nucleotides and identifying where functional components are located is complicated. With a growing number of whole genomes to work with, annotating the ORFs is more important than ever. ORFs are transcribed regions of DNA, usually genes. Without considering introns, the only criteria for a potential ORF is a start and stop codon in the same frame, meaning a multiple of three nucleotides separates them. The second goal of this project is to calculate all of the potential ORFs and their corresponding CUFs to compare with reference frequencies of the same organism to reject or deem “plausible.”

A principle concept for this ORF detection is that different organisms prefer specific codons over others. For example, out of the six codons that encode serine the human genome prefers AGC. This is relevant during translation because tRNAs for redundant codons can exist at different concentrations. Therefore redundant codons with high or low respective tRNA concentrations can either halt or speed up translation. Sometimes, this variance in translation is necessary for the growing polypeptide chain to fold correctly. So, the conventional “silent” mutations are not always silent. The main idea is that this biological quirk can be harnessed to predict correct ORFs, or rather, reject implausible ORFs. By changing the frame of a DNA sequence, the CUFs will vary greatly. If these CUFs do not align with the preferred codon patterns of the respective organism, it should be possible to reject potential ORFs. A chi-square goodness-of-fit test is used to compare the potential ORFs to a reference. The statistically significant results are rejected (the null hypothesis of the CUFs having the same distribution is rejected).

Another interesting question to ask is: Which codon frequencies are most influential on prediction? In other words, which codons account for the most variance in phylogenetic classification? Here, lasso regression and random forest feature ranking are used to rank the codons by their contribution to the classification. The overlap in the lasso and random forest results showcase the most variable codons and, therefore, possibly the most variable tRNA concentrations throughout evolutionary history. The overlap is also used separately for the statistical goodness-of-fit comparisons, which yields even better results than using all 64 codons.

### 1.2 Literature Review

This project is inspired by the work of Dr. Bohdan B. Khomtchouk, who used several machine learning approaches to predict the kingdom of a species or DNA type of a sample from its CUFs [1]. Dr. Khomtchouk used K Nearest Neighbors (KNN), Random Forests (RF), Extreme Gradient Boosting (XGBoost), Artificial Neural Networks (ANN), and Naive Bayes (NB). The kingdom results to improve upon are listed in Table 1.

**Table 1:** Highlights of the results from [1], the corresponding accuracy and macro F1-score are listed with each model.

| Model | Accuracy | F1-Score |
| --- | --- | --- |
| KNN | 96.60 | 0.9867 |
| XGBoost | 95.02 | 0.8846 |
| RF | 92.98 | 0.8611 |
| ANN | 91.32 | 0.8425 |
| NB | 75.22 | 0.7032 |

The kingdom discrimination that Dr. Khomtchouk used for their classification is detailed in the data description section below. When analyzing Dr. Khomtchouk’s results, the performance of ANN was good at 91.32%, but could definitely be improved with a more complex model. Also, an ensemble might be a natural progression to improve upon the great performance of KNN (96.60% accurate) by catching hard to classify data points.

### 1.3 Data Description

The dataset of CUFs comes from [2]. It is composed of 69 columns that list the following: 1 is Kingdom, 2 is the DNA type, 3 is a species ID, 4 is the number of total codons (Ncodons), 5 is the species name, and 6-69 are the codon usage for each of the 64 codons recorded as the number of instances divided by Ncodons. There are 13028 DNA samples total for each column. The kingdoms are categorized by three letter codes for archaea (arc), bacteria (bct), bacteriophage (phg), plasmid (plm), invertebrate (inv), vertebrate (vrt), mammal (mam), rodent (rod), primate (pri), and virus (vrl). The DNA types are listed as numbers. 0 is genomic, 1 is mitochondrial, 2 is chloroplast, 3 is cyanelle, 4 is plastid, 5 is nucleomorph, 6 is secondary endosymbiont, 7 is chromoplast, 8 is leucoplast, 9 is NA, 10 is proplastid, 11 is apicoplast, and 12 is kinetoplast.

This kingdom classification is not intuitive considering the normal (7 not 11) biological kingdom hierarchy. There is also oddly no fungi data. More extensive datasets curated from genbank and [3] may increase the performance of the models. Unfortunately, the codon usage for many organisms is surprisingly well characterized but extremely poorly organized. A significant amount of work lies ahead to compile a manageable and wide enough (phylogenetically) for future applications to be very useful. For this project the data is organized into “kingdom-set 1:” animals, plants, viruses, bacteria, and archaea by combining the relevant kingdoms (bacteriophage into virus, plasmid into bacteria, all animals into animals etc.). This set may be more relevant although slightly less discriminatory than the original paper kingdom-set, so the analysis was done on both sets for comparison.

The original dataset from [2] contained a few questionable data points; there is extra text from the species data leaking into a codon column and some missing data points. These rows are not complete sets of data, so they have been omitted from the analysis (only three total rows omitted). For analysis the data was shuffled with sklearn [4] and 20% was removed for test data.

#### 1.3.1 Test DNA for ORF Detection

The DNA used to test the pipeline was of commercially available plasmids from Addgene [5] which were picked based upon personal familiarity. Plasmids are typically small, circular components of double stranded DNA. They are usually used by bacteria to exchange genetic information, but are used commonly in molecular biology research to give bacteria genetic material. The first plasmid used is the protein expression vector PUC18 as shown in Figure 1. It is a 2686 base pair (bp) plasmid with two correct ORFs with 855 total potential ORFs. The correct ORFs are genes that encode E. coli proteins (AMPr and B-gal), so the E. coli reference frequencies will be used to compare this data. The second plasmid is for cloning HSP90 (a human protein) with an HA tag as shown in Figure 2. At 7636 bp this plasmid has 5470 potential ORFs. There are five correct ORFs in this sequence, but here just interested in the human one (HSP90), so the human reference frequencies will be used.

**Figure 1:**
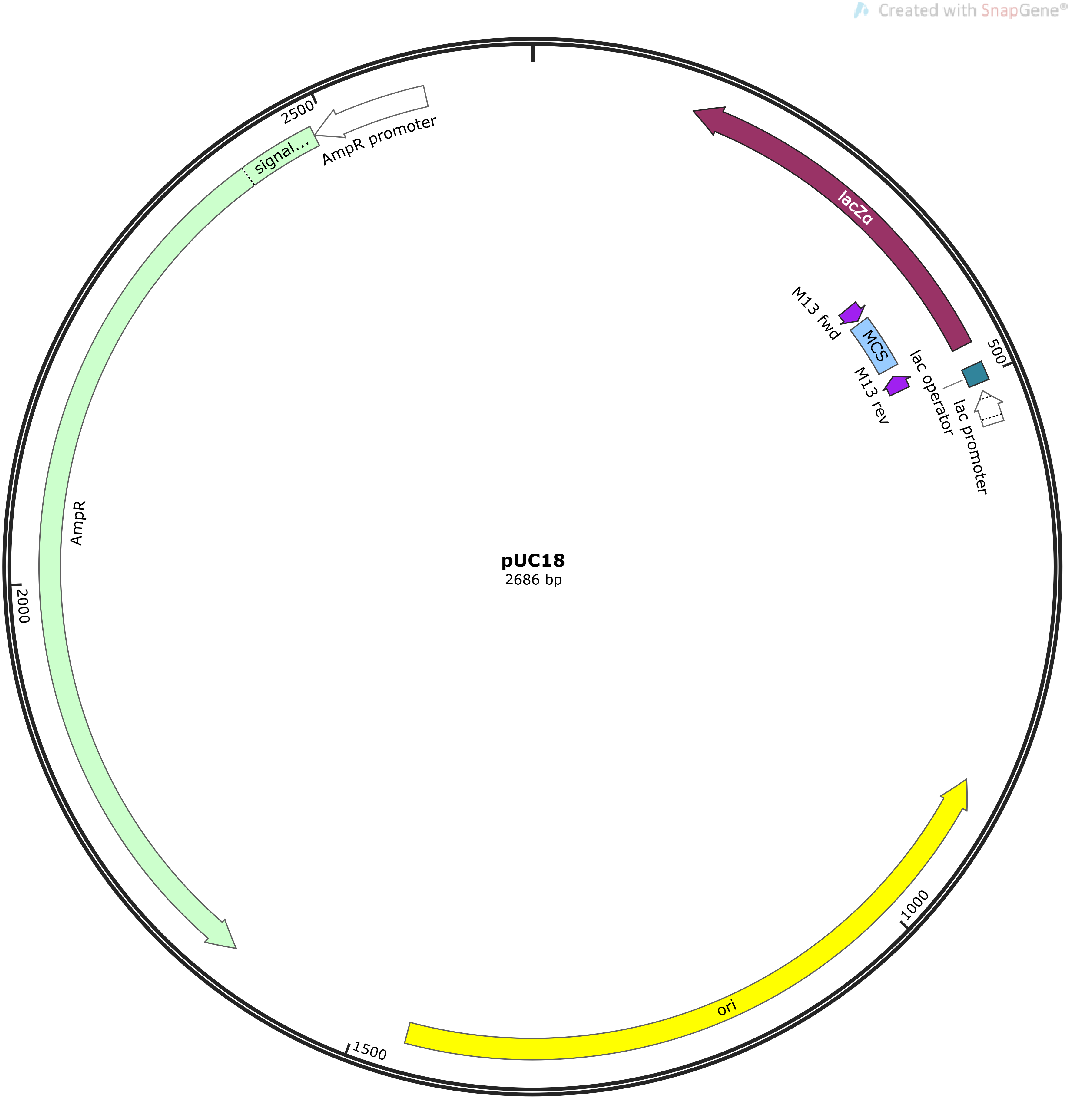
PUC18 plasmid map generated with SnapGene [6]. The 2686 bp plasmid has 855 potential ORFs with only two correct ones. They code for an ampicillin resistance protein and beta-galactosidase. This plasmid is typically transformed in E. coli and contains E. coli proteins, so a E. coli reference sequence will be used for analyzing this sequence. Some other features of the plasmid and their directionality on the DNA stands are also highlighted.

**Figure 2:**
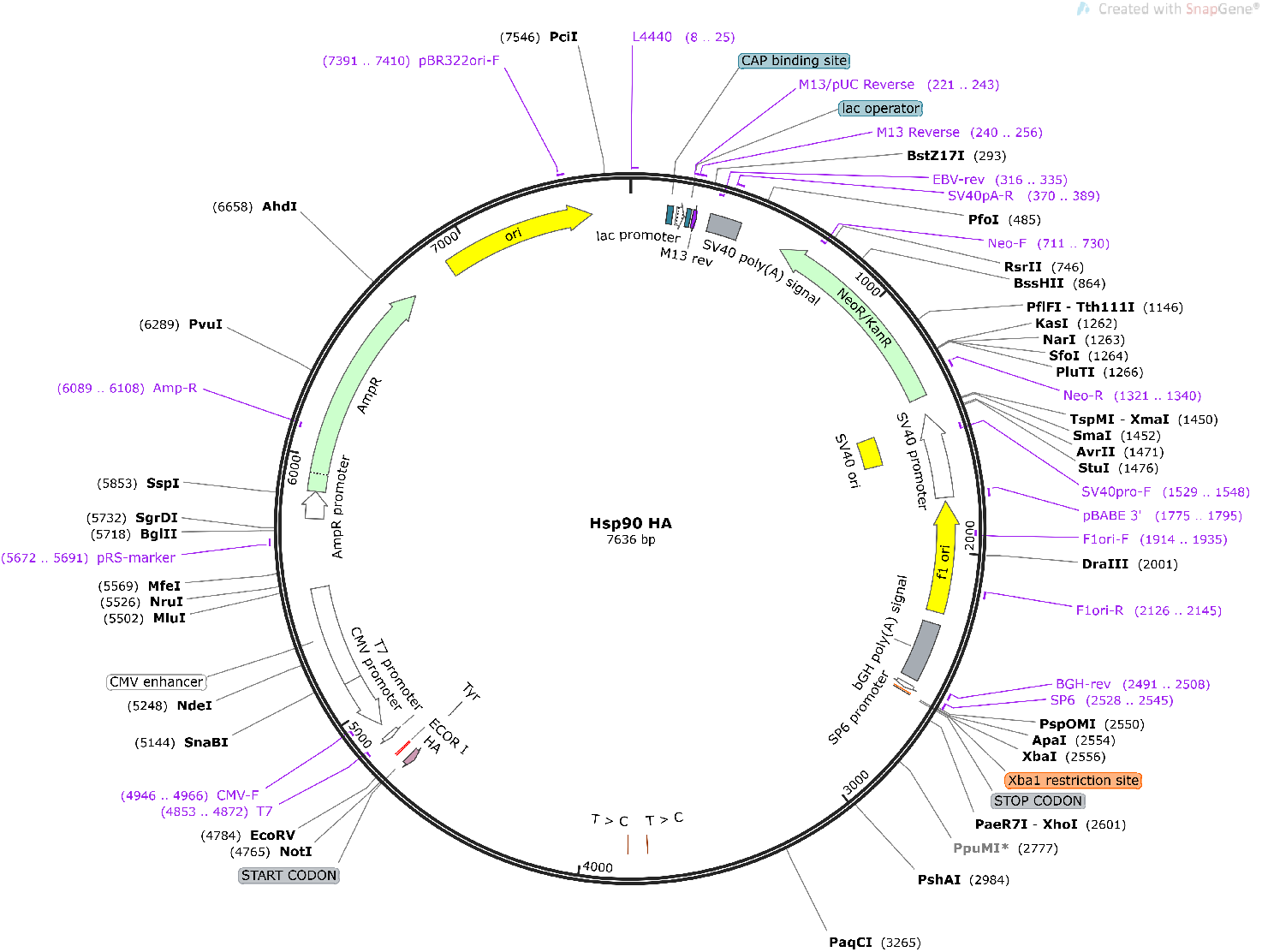
HSP90-HA plasmid map generated with SnapGene [6]. The 7636 bp plasmid has 5470 potential ORFs with only four correct ones. Here the interest is of the HSP90 ORF. This ORF codes for Heat shock protein 90 (HSP90) which is a human protein involved in the heat shock response. This plasmid is typically transformed in E. coli, and contains E. coli proteins and a human protein. Because the analysis will be done to deem the HSP90 ORF plausible the human reference sequence will be used for analyzing this sequence. Some other features of the plasmid and their directionality on the DNA stands are also highlighted.

The other sequences of DNA subject to analysis were of microsattelites and other non-coding repeats. For example, the sequence AAUGG or CACACA repeated 50 or so times resemble repeats encountered frequently in whole genome sequencing. Because genomes are scattered with simple non-coding repeats, it is important that a ORF detection tool rejects all or almost all of the potential ORFs in these regions.

## 2 Methods

All models were coded and run in python. The sklearn [4] and TensorFlow [7] packages are incredibly useful for generating these models with minimal lines of code.

### 2.1 Artificial Neural Networks

ANNs are designed to model a set of interconnected biological neurons. To simplify things, it is common to have an input layer that the data feeds to, this outputs to a certain amount of middle layers, and finally there is an output layer considering each possible label. The outputs of neurons are weighted, and if the sum of a weighted inputs meet the threshold of their activation function they “fire” outputting to the next layer. These weights are what the chosen loss function optimizes over. This project uses a full feed forward neural network, meaning that each neuron outputs to every neuron in the next layer without backpropagation. To maintain a better model and a flexible code depending on the discrimination required, GridSearchCV was used to search over hyperparameters. Using the sparse categorical crossentropy loss function, the adam optimizer, with 5 fold cross validation (CV), 15 epochs each model, and a validation split of 0.1 the code chose the number of layers, neurons in each layer, and the l2 penalty term for regularization. The code is as follows:

~~~
*#Function to b u i l d each model*
**def** build model (n_layers, n_units, l2_pen, input shape = [64]) :
 RegularizedDense = partial (Dense, activation=’relu’,
   kernel in itialize r=’he normal’,
   kernel regularize r=keras.regularizers.l2 (l2=l2_pen))
 model = Sequential ()
 model.add (layers.InputLayer (input shape=input shape))
 **for** layer **in range** (n_layers) :
   model.add (RegularizedDense (n_units))
 model.add (RegularizedDense (n_units / 2))
 model.add (RegularizedDense (n_units / 4))
 model.add (Dense (5, activation=’softmax’))
 model.**compile** (loss=’sparse_categorical_crossentropy’, optimizer=’adam’,
 metrics =[’accuracy’])
 **return** model
keras_class = tf.keras.wrappers.scikit_learn.Keras Classifier (build model)
*#Possible parameters*
params ann = *{*”n_layers” : [3, 4, 5, 6],
 “n_units” : [128, 320, 640],
 “l2_pen” : [0, 0.00001, 0.000001, 0.000001, 0.0000001]
 }
*#CV on parameters*
grid_ann = GridSearchCV (keras_class, params ann, cv = 5)
grid_ann.fit (X_train, y_train, epochs = 15, validation split = 0.1)
*#Save the best model as model*
best_neural_model = grid_ann.best_estimator_. model
~~~

Not only does this code allow for choosing hyperparameters but the final two layers are half and a quarter of the chosen number of units, allowing for a gradual decrease in node count close to the output. This is common in many deep neural networks. The range of hyperparameters chosen was narrowed down after many iterations of this code, but ultimately the choices are limited by the computation time for such a large model.

### 2.2 Ensembles

Ensembles are compositions of multiple models. Features are independently input to each model individually, and (in the model used for this project) the outputs are weighted evenly towards a majority vote for the ensemble prediction. For the individual models below, their hyperparameters were chosen using GridSearchCV with 3 fold CV.

#### 2.2.1 K Nearest Neighbors

KNN models group the *n* by *p* data matrix into *p*-dimensional feature space. It uses the labels of the *k* nearest points (in this case by euclidean distance) to classify the datapoint of choice. *k* = 1 performs best for kingdom-set 1.

#### 2.2.2 Support Vector Machines

Support vector machines (SVM) scale up the dimensions of the feature data until it can be separated by a function. The best hyperparameters for kingdom-set 1 are *C* = 11, *γ* = 188, and using an rbf kernel.

#### 2.2.3 Random Forests

RF models are an ensemble method themselves as the composition of many decision trees. Each tree makes a prediction based on the labels of the feature data, the predictors are randomly cho- sen. The hyperparamaters of choice are n estimators=533, max features=sqrt, max depth=none, min samples split=2, min samples leaf=1, and bootstrap=False.

The RF model was also used to rank the codons in terms of importance for prediction. RFs can do this by keeping track of how each split and feature improve the prediction from the training dataset. The higher the leaf purity, the higher the importance of the feature. When this is kept track of in each tree for all trees, it can be normalized to 1 and the output is the “importance score” of each feature [8].

#### 2.2.4 Others

Logistic regression, naive bayes, and another decision tree models were also trained and optimized for the data. However, when they were added to the ensemble they decreased the test accuracy vs. just KNN, SVM, and RF. Therefore, they were not kept in the final model. The final model is as follows:

~~~
knn_clf = KNeighborsClassifier (n_neighbors = 1)
rf_clf = RandomForestClassifier (n_estimators = 533, min_samples_split = 2,
  min_samples_leaf = 1, max_features = ‘sqrt’, max_depth = None, bootstrap = False)
svc_clf = SVC(C = 11, gamma = 188, kernel = ‘rbf’)
voting_clf = sklearn.ensemble.VotingClassifier(
  estimators =[(’knn’, knn_clf), (’rf’, rf_clf), (’svc’, svc_clf),], voting=’ hard ’,)
~~~

### 2.3 Lasso Regression

Lasso regression is a penalized regression method that adds to the loss function when a coefficient is added to a parameter with an l1 norm.

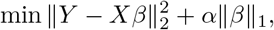

Where the feature data *X* times the “learned” coefficents *β* estimate the label vector *Y*. *Y* = *Xβ* is the absolute minimum for linear least squares regression which means a perfect estimation, however, with the penalty constant *α* times *β* the coefficients must be small and estimate *Y* well. Therefore, as the penalty constant gets bigger, the features must be very good predictors if it is going to be worth adding them to the model. This allows trimming and ranking features (in this case codon frequencies) by varying the penalty. Lasso is a simple convex optimization problem that most computers can solve extremely quickly, so it is easy to set up a loop that trains codon frequencies vs. kingdom-set 1 and varies *α*. A good loop will start extremely small and change a tiny amount each time in order to track the order of each feature leaving the coefficients. What worked for this data is started at *α* = 10^*−*8^ and varying by 10^*−*6^ until *α* = 0.01 is reached. The code allows tracking of the coefficients in front of each codon, and then take the 0 norm of the column vector to count how many nonzero entries there are. Then, the most nonzero entries account for the most variance in the prediction and are ranked accordingly.

### 2.4 K Means

K Means is a clustering algorithm that optimizes *k* centroids to separate the feature data. It does not use the labels, so it is an unsupervised model. The idea behind using this with vast *k* values is to see if there are any higher *k* models that separate the data well. kingdom-set 1 is similar to *k* = 5, because there are 5 categories. The original kingdom-set is approximately equivalent to *k* = 11. If the codon usage data is well separated by another more discriminatory structure, perhaps biological order, that might show up in K Means analaysis.

#### 2.4.1 Potential ORF Codon Frequencies

The potential ORFs of a sequence are calculated by a class in python. The class is setup to take an input string of RNA nucleotides A, G, C, and U. DNA sequences are easily converted to RNA sequences by replacing all T’s with U’s. The initial function finds all start codons (AUG) and all stop codons (UGA, UAG, UAA) and stores the index of the first nucleotide of them all. If the start and stop are separated by a multiple of three nucleotides, and the index of the stop is greater than the index of the start, this region is recorded as a potential ORF. The potential ORF sequences are all fed to an additional function that reads the codons in frame and records the frequency of each one by dividing the number of instances by the total number in the sequence. All potential ORF frequencies are stored in a three dimensional array that is easily searchable by size using the len() function. This is useful for looking for expected ORFs and it was easy to find the correct ORF frequencies from the plasmid sequences discussed in the data description section.

It is important to note that this class only reads the sequence in one direction, and because DNA is double stranded ORFs can exist reading 5’ to 3’ or 3’ to 5’ (the directionality of DNA). This is okay for testing known DNA sequences as a proof-of-concept, but for high throughput and thorough analysis, future iterations of the code should include a function that calculates the reverse compliment and sends that through the class as well. There should also be a function that tests for DNA and converts to RNA for ease of use (or vice versa, having DNA or RNA is not important but consistency throughout is). The program MegaX [9] can do both of these easily, which is what was used to help with the preliminary analysis.

#### 2.4.2 Goodness of Fit Tests

The chi-square goodness of fit test is used to statistically evaluate if a set a sample “frequencies” come from a population “frequency” of a specific distribution [10]. This test is defined for the null hypothesis: The sample data follows a specified population distribution. It is advantageous for the purposes of testing ORF frequencies because it is extremely flexible on binned data which ORF frequencies are somewhat analogous to. The disadvantage of this method is that it requires a sufficient sample size, the standard being 5 or larger in each bin. Frequency is in quotations above because chi-square really requires a count, which is easy to calculate by multiplying the frequencies by the total number of codons in the potential ORF. The population frequency from the organism of interest is also multiplied by the same number for consistency.

For the computation, the data separated into *k* bins (64 for codon usage) defines the test statistic as

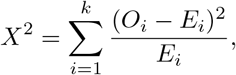

where *O*_*i*_ is the observed count for bin *i* and *E*_*i*_ is the expected count (from the population) for bin *i*. In the case of CUFs we use *k −* 1 degrees of freedom assuming that the distributions are roughly multinomial. With this in mind, the null hypothesis is rejected (a non-plausible ORF) when

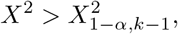

where 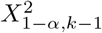 is the chi-square critical value using *k −* 1 degrees of freedom and a significance *α*. The typical standard for *α* is 0.05; however, in this application the sample frequencies are extremely easily rejected and tuning *α* for useful output is tricky. Importantly, none of the population counts can be 0 due to the *E*_*i*_ term in the denominator. 0 counts are not found in every species from [2] but are not uncommon either. Usually, there is only one 0 count in the reference if any, so it is not a huge issue. To accommodate, the python class looks for any 0s in the population frequency and the corresponding codons are removed from the sample and population frequencies. The chi-square goodness of fit test is implemented in python under scipy.stats, and automatically calculates the degrees of freedom during calculation; thus this removal of codons does not significantly change the results of the test.

The Cressie-Read goodness of fit test was also tried for rejecting potential ORFs. The Cressie-Read test is similar to the chi-square test but it can account for 0s in the population count [11]. The difference in the test statistic and *p*-value when comparing this adjusted chi-square vs. the Cressie-Read was negligible. Over several samples chi-square with degree of freedom 62 for 63 codons vs. Cressie-Read for 64 codons performed almost exactly the same.

## 3 Results

### 3.1 Artificial Neural Network Performance

The ANN model from [1] was not sufficiently complex and hyperparamters do not appear to be optimized, resulting in an accuracy of 91.32%.

The best performing model for kingdom-set 1 performed much better. It was trained with 20 epochs resulting in 95.55% accuracy and 0.9302 macro-F1 score on kingdom-set 1. The model gave a 93.55% accuracy and 0.8627 macro-F1 on the original kingdom-set. The F1 score is 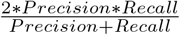, meaning that a perfect model has a score of 1. The confusion matrix for this model is highlighted in Table 2.

**Table 2:** Confusion matrix for optimized ANN model kingdom-set 1. The entries on the left to right diagonal represent correctly classified test data points Ex. virus as virus. The others are misclassified, Ex. plant as animal.

|  | Virus | Bacteria | Archaea | Plant | Animal |
| --- | --- | --- | --- | --- | --- |
| Virus | 562 | 32 | 1 | 14 | 14 |
| Bacteria | 1 | 573 | 3 | 7 | 0 |
| Archaea | 0 | 2 | 24 | 0 | 0 |
| Plant | 4 | 11 | 0 | 484 | 13 |
| Animal | 8 | 13 | 2 | 11 | 827 |

### 3.2 Ensemble Performance

The ensemble of optimized KNN, RF, and SVM performed even better than ANN on test data with an accuracy of 97.16% and macro-F1 of 0.9562. The original kingdom-set yielded an accuracy of 99.08% and macro-F1 of 0.9781. The confusion matrix for this model is highlighted in Table 3.

**Table 3:** Confusion matrix for the final ensemble on kingdom-set 1. The entries on the left to right diagonal represent correctly classified test data points Ex. virus as virus. The others are misclassified, Ex. plant as animal.

|  | Virus | Bacteria | Archaea | Plant | Animal |
| --- | --- | --- | --- | --- | --- |
| Virus | 600 | 15 | 0 | 4 | 4 |
| Bacteria | 4 | 572 | 2 | 5 | 1 |
| Archaea | 0 | 2 | 23 | 1 | 0 |
| Plant | 5 | 4 | 0 | 494 | 9 |
| Animal | 12 | 1 | 0 | 9 | 839 |

### 3.3 Ranking Codons

Both feature rankings are highlighted in Figure 3.

**Figure 3:**
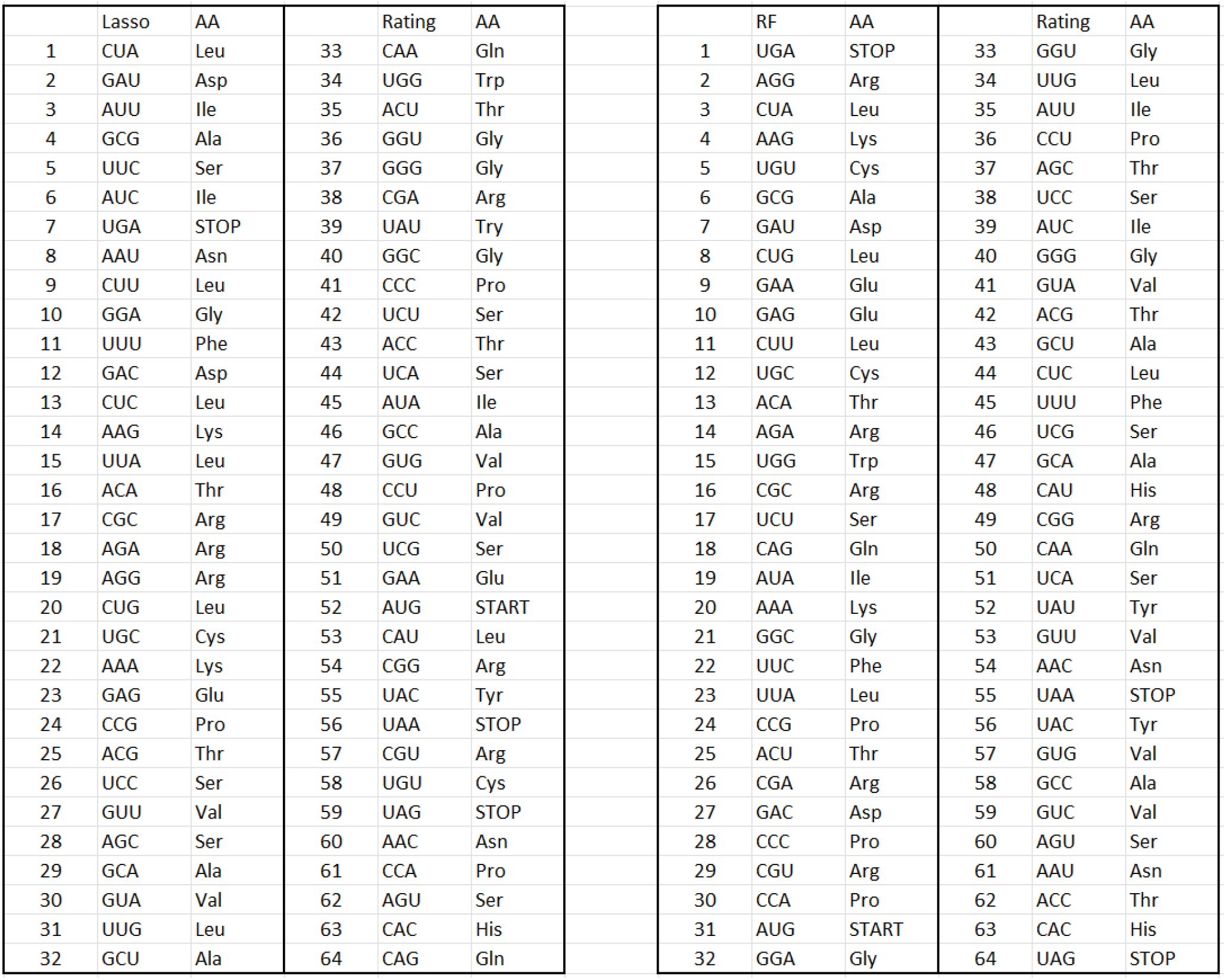
Sorting CUFs and their corresponding amino acid by their prediction power with lasso and RF.

Here the lasso and RF ranking choose a different feature order but with a clear preference to the order of certain codons. Regardless of the discrimination chosen (euk vs. pro or more specific) CUA comes up as the most influential from the lasso analysis. The codons in the top 20 of both ranking systems is CUA, GAU, GCG, UGA, CUU, AAG, ACA, CGC, AGA, and CUG. Perhaps this is a starting place for uncovering novel biological function in tRNA concentration and usage.

### 3.4 Clustering

K Means analysis was conducted on the set of 13026 codon frequencies for clusters 2-600. An elbow graph and corresponding silhouette scores for clusters 2-50 are shown in Figure 4. The goal here was to uncover any unintuitive clustering structure to gain insight into how specific the codon usage phylogenetic classification would go. The hope was that the silhouette score would jump up or an interesting pattern in inertia would appear, indicating the possibility of a larger *k* clustering in the data. This was not the case past clusters 35-39 with a large jump in silhouette and slight increase in inertia. This may indicate some predictive ability to classify on the order of phylum, class, or order but certainly not species.

**Figure 4:**
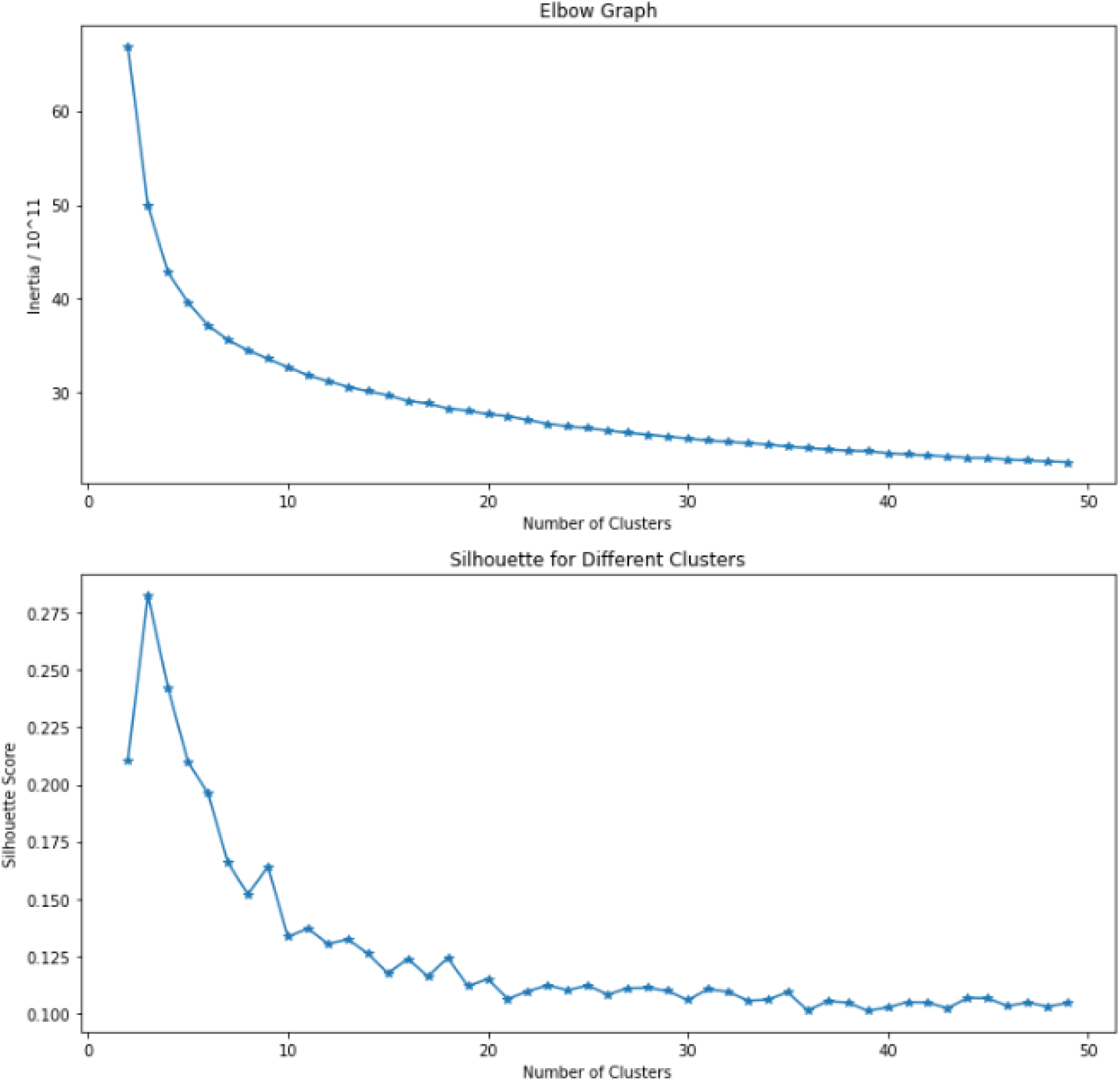
Elbow graph and silhouette score by cluster number generated from K Means on CUF data. Listed are clusters 2 to 50. The inertia and silhouette pattern highlight the most intuitive clustering pattern at *k* = 3 with decent values for our kingdom-sets *k* = 5 (analogous to kingdom-set 1) and *k* = 11 (analogous to the original kingdom-set).

#### 3.4.1 ORF Detection

The sequences input to the python class gave some interesting results after the statistical test. When comparing all 64 frequencies to an E. coli reference for PUC18 all 855 potential ORFs are rejected with tiny *p*-values. The same occurs to all 5470 potential ORFs of the HSP90 plasmid using a human reference. However, the *p*-values for the correct ORFs are much bigger than their surrounding incorrect ORFs; correct hovering around *p* = 10^*−*50^ while incorrects vary from 10^*−*100^ to 10^*−*200^. Therefore, the test is much more confident that the incorrect ORFs were incorrect compared to the correct ones. This hints that there is possible for optimizing in specific applications of this method to more likely reject incorrect ORFs.

The results are much more reasonable when only using the most influential codons listed above (the shared codons in the top 20 of lasso and RF analysis). Using a *p*-values threshold of 0.05 only 9 ORFs are rejected from the HSP90 plasmid and 12 are rejected from PUC18. More importantly, none of the correct ORFs from either were rejected. This is much more in line with the goals of the project: creating a tool in addition to well studied methods of ORF detection to trim a small amount of the potential ORFs.

When tried on microsatellites and tandem repeats common in human genome annotation the statistical test unsurprisingly rejects all potential ORFs on the order of *p* = 10^*−*300^ or even 0 output. This is promising because the statistical test should reject these repeats extremely confidently, as the whole idea behind the predictive power here is that if certain codon frequencies get too high, it is unlikely to be a correct ORF. With a huge sequence of repeated nucleotides, the codons will also be repeated and luckily the test gives the expected results.

## 4 Discussion

After careful selection of hyperparameters with cross-validation and consideration for factors such as overfitting, under / oversampling, and more the newer models clearly perform better than the ones referenced from the literature. GridSearchCV proved to be a valuable tool for setting up many hyperparameters with many models and then optimizing overnight. For instance, with minimal lines of code GridSearchCV was able to test all the hyperparameters over a decent range for KNN, RF, decision trees, multinomial naive bayes, logistic regression, and SVM with three-fold cross-validation at the same time. It also allowed something similar for ANN by building a function that varies the number of layers, number of neurons, and an l2 penalty to prevent overfitting.

Kingdom-set 1 was designed to be more intuitive, lose minimal discrimination, and even out the categories for consideration about under / oversampling. For both kingdom-sets, the ensemble performed better than the ANN model, with an incredible 99% test accuracy for the ensemble on the original kingdom-set. This trend might fall towards ANN if more data is added to the dataset, as ANNs can better learn the attributes of the data when there is more of it. Interestingly, the ANN got worse comparing kingdom-set 1 to the original, but the ensemble got better. This may be because of an ensemble’s ability to classify tricky data points, even when one individual model would miss it. The relevant results are shown in Table 4.

**Table 4:** Comparison of original ANN model from [1] with the improved models over two different kingdom-sets: Original and Kingdom-set 1.

|  | Original Set |  | Kingdom-set 1 |  |
| --- | --- | --- | --- | --- |
| Model | Accuracy | F1-score | Accuracy | F1-score |
| Original ANN | 91.32 | 0.8425 | NA | NA |
| Improved ANN | 93.55 | 0.8627 | 95.55 | 0.9302 |
| Improved Ensemble | 99.08 | 0.9781 | 97.16 | 0.9562 |

The main future work for broadening the capability and applications of this classification is adding more data. There are vast datasets (50+ Gb excel files!) full of all genbank [3] entries with CUFs. However, this amount of data is not manageable with current household computer machinery. Hopefully, future capabilities allow the extraction of specific new data (especially fungi).

Overall, this ORF detection (or rejection) pipeline establishes a basis for a small amount of predictive power between CUFs and ORFs. The results are most promising when comparing the most influential codon frequencies found with lasso and RF feature ranking. However, this ranking and comparison has a huge amount of variability and room for improvement. The totally arbitrary choices in the selection of which codons include the lowest common rank (20) and the discrimination used for phylogenetic classification (kingdom-set 1). Different discrimination or the lowest common rank, for example, biological phylum and 30, might give even better or worse results. It may also be helpful to use the most preferred redundant codons from an organism for annotation in that organism. For instance, there are six codons that code for serine: UCU, UCC, UCA, UCG, AGU, and AGC. Only one of these codons is most common in each organism, so choosing the most common redundant codon may have more predictive power on an organism by organism basis. Another level of complexity is choosing by which codons are matched to which tRNA. For example, there is potentially one tRNA molecule for UCU, UCC, UCA, and UCG (the last nucleotide is known as a “wobble” nucleotide) and one tRNA molecules for AGU and AGC. Therefore, choosing the preferred tRNA set of codons is another possibility for optimizing this method.

Clearly, there is a lot of work to be done to optimize this method. On top of choosing which codons to test the *α* for useful rejection is unknown. An entirely different statistical test may be necessary as well. Choosing an *α* may be possible with some regression or classification methods on a labeled data set composed of half correct ORFs and incorrect ORFs. While this may seem promising (and tedious), it would probably only optimize the method for a specific organism or phylum at best. It may need to be redone for each organism being annotated. Choosing *α* this way appears to only be necessary when considering all 64 CUFs. The vast complexity of the distribution in 64-dimensional space means that none of the potential ORFs are anywhere near close enough to be considered the same distribution. This is to be expected, each protein has a varied amino acid sequence that should not align with the average sequence of the entire organism. It is only by looking at particularly varied or influential codons that there can be predictive power here on the order of *α* = 0.05 or 0.01.

It would be fascinating to take this further and implement it at the end of large-scale genome annotation pipelines. This is almost necessary for the method to be helpful because of one massive problem: introns. Introns are the sections of mRNA that get spliced out of primary mRNA sequences in order to form the final mRNA product, composed of the actual coding regions. These coding regions are called exons. The CUFs from ORFs all come from exons, meaning that the intron sequences of DNA (non-coding regions in the middle of ORFs) may completely ruin this method. Luckily, biologists have discovered many properties of introns that make them possible to identify. For example, the splice sites of introns are usually the sequence AG—GU and AG—G (— is where the sequence is cut). Therefore, when they are spliced and put back together the sequence continues AGG like there was nothing there in the first place. Because of these intron properties (and more), there are well-studied modeling techniques that can evaluate splice sites and identify possible introns, potentially making CUFs a useful high throughput method. There are also other functional non-coding sequences of DNA that could also be leveraged. For instance, if a start codon is near a promoter sequence, it should be leveraged as much more likely to be correct than if not.

Thus, with all of these considerations at play and optimization to be done, there is decent potential in this method to produce useful results. Biology is complicated because of the complexity and stochasticity at every step; however, it can become more manageable with careful mathematical modeling. As CUF phylogenetic classification improves and datasets become larger and more vast in biological depth, ORF detection and more applications should come to the surface using this neat quirk of biological regulation.

## Acknowledgments

This project is the culmination of three final projects for graduate classes at the University of Delaware, and so I would like to acknowledge the crucial advice and guidance I received from the following professors: Dominique Guillot, Ph.D., Mark Keese, Ph.D., and Li Liao, Ph.D.

